# Genome-mining algorithm to identify identical repetitive sequences for sensitive and specific diagnostic assays for infectious diseases

**DOI:** 10.1101/2024.12.20.629856

**Authors:** Kalepu Rajeswari, Raksha Poojary, Shruptha Padiwal, Rajesh Muliyar Krishna, Kapaettu Satyamoorthy, Bobby Paul

## Abstract

Nucleic acid amplification-based approaches are extensively used as the first line of choice for infectious diseases. However, the success rates of DNA amplification or hybridization techniques are highly dependent on short primer or probe sequences. A pair of primers that can bind at multiple loci across the genome and randomly amplify multiple copies increases the analytical sensitivity of the currently used diagnostic assays. Herein, we developed a novel genome mining algorithm to identify short identical repeat sequences (IRSs) dispersed across the genome, which can amplify multiple nonhomologous regions of variable sizes via three potential priming combinations. Using this algorithm, we analysed the genomes of five pathogens, namely, gammaherpesvirus, vaccinia virus, *Mycobacterium tuberculosis, Plasmodium falciparum*, and *Phytophthora palmivora*, and identified short identical sequences that were repeated at multiple loci. *In silico* PCR revealed that these identical repeat sequences can amplify multiple copies with different amplicon sizes in these five species. We further performed a polymerase chain reaction assay with short identical repeat pairs identified from *M. tuberculosis*. Very interestingly, the amplification yielded multiple copies for individual IRSs and even more copies, as in a pair of IRSs. These results indicate that the IRS-based approach can detect pathogens during disease progression in the case of low-concentration DNA. The genome mining algorithm can be used as a translation technology platform for developing highly sensitive varieties of PCR, microarray, loop-mediated isothermal amplification, fluorescence *in situ* hybridization, and DNA-DNA hybridization-based diagnostic assays.

## Introduction

Infectious diseases are perpetual public health concerns, and new infections may continue to appear in the future (Piret and Boivin, 2021). Quick, low-cost, and accurate identification of pathogens is essential for diagnosing and treating infectious diseases. Over the years, microscopic imaging to high-throughput multiomics approaches have evolved to identify pathogens accurately (Franco-Duarte et al., 2019; Gerace et al., 2022). Nucleic acid-based assays are superior to most currently available infectious disease diagnostic approaches because i) almost all pathogens carry nucleic acids, ii) DNA has very good potential to serve as a biomarker, and iii) these can increase the sensitivity and speed of detection during the early stages of disease (Chen et al., 2002; Chen et al., 2019; Lam et al., 2020). As a culture-independent approach, isothermal-based nucleic acid amplification and detection assays are cost-effective, rapid, highly sensitive, and user friendly (Niemz et al., 2011). Furthermore, nucleic acid detection (NAD)-based real-time polymerase chain reaction is considered the gold standard for infectious disease diagnosis (Fang et al., 2023).

Deoxyribonucleic acid (DNA)-based tests are extremely important and are extensively used in hybridization, thermal-cycle or isothermal or signal amplification, and microfluidic assays for pathogen detection (Li et al., 2021). Primers and probes are essential components for DNA-based diagnostic assays involving amplification or hybridization. Hence, we developed a novel genome mining algorithm to identify pairs of short identical sequences that constitute the maximum number of repeats in a genome and explore their potential as primers or probes. Several studies have reported that the use of multicopy genes may increase the detection sensitivity of DNA-based assays. A highly specific and sensitive loop-mediated isothermal amplification assay targeting the multicopy gene *ORF160b* was developed for the detection of allergic soybeans (Allgower et al., 2020). A few other studies have reported that multicopy gene-targeted DNA-based assays improve the detection sensitivity of malarial parasites (Hofmann et al., 2015; Gupta et al., 2016; Nolasco et al., 2021).

A study was performed with a genome mining approach to identify a pair of identical multirepeat sequences that constitute multiple copies in the *Plasmodium falciparum* genome and explore their potential as primers for PCR (Raju et al., 2019). A recent study also identified identical multirepeat sequences as biomarkers to diagnose *N. gonorrhoeae* (Shiluli et al., 2024). However, these algorithms require genome fragmentation with a defined length (L-mer) in all possible combinations to identify identical multirepeat sequences. Advancements in next-generation sequencing technologies have increased the availability of large amounts of genomic data from different pathogens. There is an urgent need to develop algorithms to translate genomic data to DNA-based technologies that are widely acceptable for small- or medium-sized laboratories (Church et al. 2020). Therefore, we sought to develop a new genome mining algorithm to identify identical repeat sequences (IRSs) that randomly amplify and constitute a large number of fragments from the genomes of pathogens. The results, such as effort, are presented.

## Methodology

### Genome mining algorithm development

The algorithm for IRS identification is written in the Python scripting language, as it is ideal for prototype applications and has excellent text processing capabilities. The algorithm searches for the number of occurrences of user-defined length (k-mer) sequence patterns in forward and reverse complement sequences in a DNA strand, as illustrated in Figure 1. Furthermore, the sequence patterns will be sorted in descending order on the basis of their repeat count. The user can provide the input values for three parameters before executing the algorithm, and the value must be a number. The first parameter is the k-mer length: the user can define the k-mer length, which is optimal for the diagnostic assay (a number between 18 and 24 is ideal for primers). The second parameter is the repeat count: the algorithm searches for identical sequences present in the given genome sequence equal to or above the indicated number. This parameter value can be comparatively small, ranging from 5-100 for organisms with a genome is small in size (up to 20 Mb). The value can exceed 100 for an organism with a genome size is greater than 20 Mb. Last, the GC content: patterns with low GC content can be eliminated with the option provided. Patterns with 50–60% GC content is ideal for PCR-based assays. As illustrated in Figure 1, a pair of IRSs can amplify genomic regions in three possible combinations. Potential pairs of IRSs that can amplify multiple regions have great potential to translate as biomarkers in DNA amplification or hybridization reactions.

**Figure 1.**
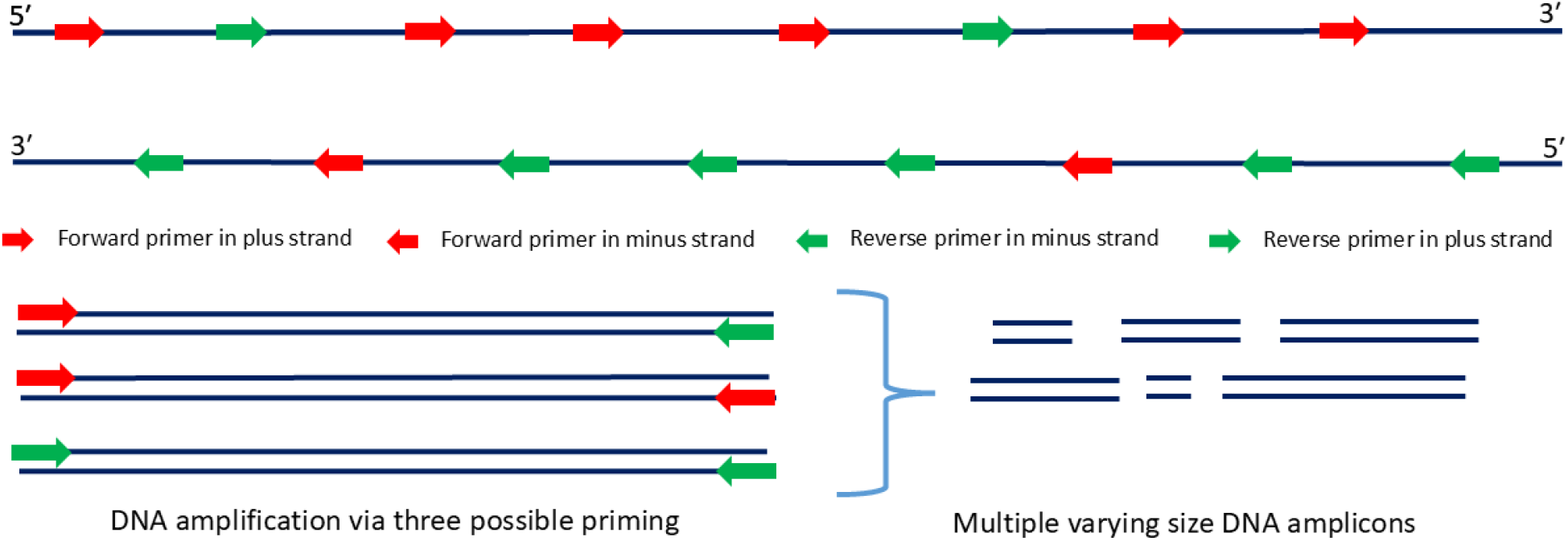
Nonhomologous DNA amplification via identical repetitive sequences enables the generation of multiple amplicons of varying sizes.

### Genome mining algorithm validation

This study used five categories of pathogens for *in silico* validation: i) human papillomavirus, *which* can cause cervical cancer; ii) vaccinia virus, which belongs to the poxvirus family; iii) *Mycobacterium tuberculosis*, a potent human pathogen that causes tuberculosis; iv) *Plasmodium falciparum*, which can cause malarial infection; and v) *Phytophthora palmivora*, that cause bud and fruit rot on many plantation crops. The whole-genome sequences of these pathogens were retrieved from the Genome database (https://www.ncbi.nlm.nih.gov/home/genomes/), and their descriptions are listed in Table 1. Among these five pathogens, *human gammaherpesvirus* 4 has the smallest genome size (171.8 kb), and *Phytophthora palmivora* has a relatively large genome size of 165.6 Mb. The G+C content is an important parameter for selecting an ideal primer/probe for DNA amplification or hybridization purposes. Hence, we selected genomes with high and low GC contents for algorithm validation. *Plasmodium falciparum* had the genome with the lowest G+C content (19.5%), and *Mycobacterium tuberculosis* was GC-rich (65.5%) and was used for IRS identification. The developed genome mining algorithm was executed in the Linux operating system with these five whole-genome sequences separately.

**Table 1.** The list of pathogens used for the *in-silico* screening of the genome mining algorithm for IRS detection.

| Sl. No. | Organism | Genome size | GC content | Accession No. |
| --- | --- | --- | --- | --- |
| 1 | *Human gammaherpesvirus 4* | 171.8 kb | 59.50% | NC_007605.1 |
| 2 | *Vaccinia virus* | 187.2 kb | 33.15% | OR515687.1 |
| 3 | *Mycobacterium tuberculosis* | 4.41 Mb | 65.50% | CP130807.1 |
| 4 | *Plasmodium falciparum* | 23.3 Mb | 19.50% | GCF_000002765.6 |
| 5 | *Phytophthora palmivora* | 165.6 Mb | 49.93% | GCA_008079305.1 |

### Genome data preprocessing

Assembled genome sequences may not be in a uniform format. Many organisms have more than one chromosome or extrachromosomal genetic material. Furthermore, genomes assembled in the draft stage contain multiple fasta files. Hence, it is important to preprocess genome sequence data for efficient mining of identical short repeats. Shell scripts were used to preprocess the genome sequence data. The scripts convert genome sequences that contain lowercase letters to uppercase letters and remove line breaks and white spaces. Additionally, the script merges multiple fasta sequences in a file and creates a ready-to-use whole genome in the preferred file name.

### In silico PCR

Computational virtual PCR can assist in the selection of potential pairs of primers and in identifying the expected amplicons before any wet-lab experiment. Primer-BLAST (Ye et al., 2012), a web-based *in silico* PCR tool, was used to select the best IRS with primer potential. The IRSs identified from the five pathogens in the study were used as primers for *in silico* PCR. The database selected ‘custom’ with the respective accession numbers, the organism field was ‘empty,’ and primer specificity stringency parameters were set as 1 while running the program. The remaining parameters were retained as defaults.

### PCR assay for *Mycobacterium tuberculosis*

Total DNA from *M. tuberculosis* grown in culture medium was extracted via the phenol-chloroform protocol. PCR was performed in a final volume of 25 μL containing 5.0 μL of buffer (10x), 2.0 μL of MgCl_2_ (50 mM), 0.2 μL of dNTPs (25 mM), 25 pmol of each oligonucleotide (MtF1-5′ CGC CGT TGC CGC CGT TGC 3′ and MtR1-5′ GCC GCT CCT CCT CAT CGC 3′) and 0.5 μL of Taq DNA Polymerase (500 U) from Invitrogen®. Amplification was carried out for 40 cycles, each consisting of initial denaturation at 94°C for 2 min, denaturation at 94°C for 30 seconds, annealing at 58–62°C for 2 minutes, and extension at 72°C for 1 minute, followed by a final extension at 72°C for 10 minutes. The PCR products were separated via 1% agarose gel electrophoresis with a 100 bp DNA ladder. After one hour of separation, images of the DNA fragments on ethidium bromide-stained gels were captured via a gel documentation system from Thermo Fisher Scientific.

## Results

In the present study, we developed a new algorithm for genome-wide mining that identifies user-defined k-mer length sequence patterns distributed throughout a genome. The algorithm searches only for 100% identical sequences from the plus strand and their corresponding reverse complement from the minus strand. The algorithm will find identical sequences and result information such as the IRS count in plus and minus strands and the G+C content. The output file is sorted in descending order of the repeat count, and the fields are separated by a tab space. These short identical repeat sequences can be effectively used to develop various assays for infectious disease diagnosis. For conventional PCR, primers are designed to amplify specific genomic region(s). However, herein, this algorithm identifies identical short repeats that can act as forward and reverse primers and initiate amplification at multiple loci across the genome. Furthermore, the forward and reverse primers alone can amplify randomly and in pairs. Thus, in turn, this approach improves the overall diagnostic sensitivity by amplifying multiple amplicons with different fragment sizes. The algorithm facilitates robust data analysis, and compilation is optimized for runtime efficiency. In the current application, the algorithm performs genome mining on the assembled genomes to identify short identical repeated sequences of any length.

### Genome mining algorithm validation

The sensitivity and specificity of PCR-based diagnostic assays are highly dependent on the efficiency of the primer. A forward and reverse primer pair that can amplify genomic regions from multiple loci improves the overall diagnostic sensitivity and specificity. The algorithm identified short identical repeat sequences from the genomes of five pathogens, including human gammaherpesvirus, vaccinia virus, *Mycobacterium tuberculosis, Plasmodium falciparum*, and *Phytophthora palmivora*. The IRS algorithm was able to identify unique sequences from genome sizes varying from 171.8 kb to 165.6 Mb, and their G+C content varied from 19.5 to 65.5%. The algorithm was executed only for 18-20 nucleotide length patterns with more than 30% GC content. However, each genome’s repeat count (minimum) was set differently, as the genome size differed. As expected, the number of identical repeats was lower for organisms with small genome sizes, and the copy number increased with genome size (Table 2).

**Table 2.**
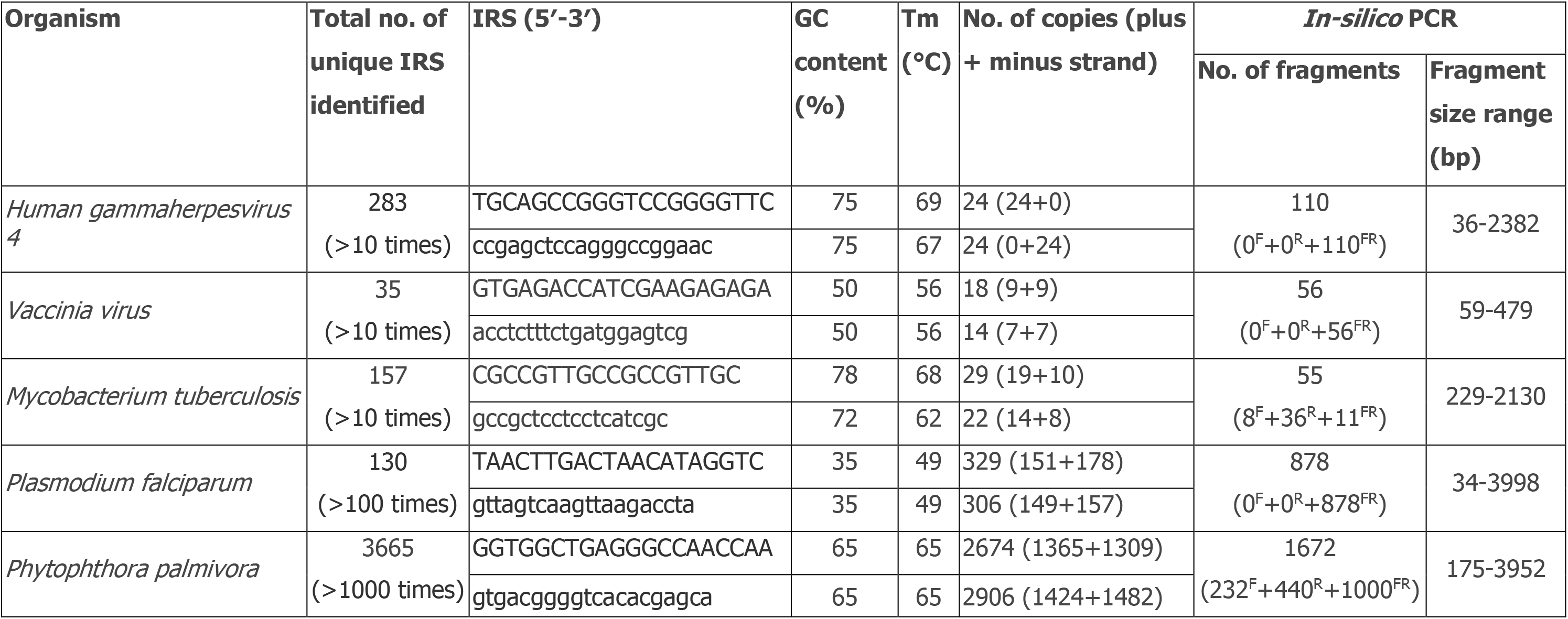
List of short-length (20 nt) IRS with good primer potential identified for PCR amplification. The table shows the best primer pair identified from each genome and their *in-silico* PCR output. Tm: melting temperature of primer. The IRSs in uppercase are considered forward, and those in lowercase are considered reverse. *In-silico* PCR was performed in three possible combinations: forward alone; R: reverse alone; FR: combination of forward and reverse.

The IRSs with high primer quality potential, as listed in Table 2, were used for *in-silico* validation, and their respective genomes are listed in Table 1. The algorithm identified a total of 283 unique identical sequences (20 nt) that were repeated more than 10 times in the genome of human gammaherpesvirus. The IRS-based primer amplified 110 fragments via *in silico* PCR, and their size varied from 36 bp to 2382 bp. A total of 35 unique identical sequences (20 nt) were repeated more than 10 times in the vaccinia virus. *In silico* PCR revealed 56 fragments with sizes ranging from 59 to 479 bp in the vaccinia genome. Similarly, 157 IRSs (20 nt) were repeated more than 10 times in the *M. tuberculosis* genome. The *in-silico* PCR analysis revealed 55 amplicons, and their size varied from 229 to 2130 bp. Furthermore, the pair of IRSs listed in Table 2 was used for *in vitro* DNA amplification of the *M. tuberculosis* genome. Furthermore, we identified another pair of IRSs (5′-GTT TCC GTC CCC TCT CGG-3′ and 5′-GTC GTC AGA CCC AAA ACC-3′), which are very specific to *M. tuberculosis* amplification. Interestingly, the pair amplified only 11 strains of *M. tuberculosis* via *in silico* PCR with entire nonredundant (nr) sequences in the NCBI database. This result highlights that the selected IRS has the potential for species-specific amplification of *M. tuberculosis*.

The algorithm identified 130 IRSs (20 nt) that were repeated more than 100 times in the genome of *P. falciparum*. The IRS 5′-AAC CCT AAA CCC TAA ACC CT-3′ was found to be repeated a maximum of 555 times (266 times in the plus strand and 289 times in the minus strand). *The in-silico* PCR results revealed that 878 amplicons were amplified from the selected IRS, and their size varied from 34 bp to 3998 bp. The genome of *P. palmivora* contains 3665 unique IRSs (20 nt) that are repeated more than 1000 times. The IRS 5′-GCA AGG TGC TCG TGT GAC CC-3′ repeated a maximum of 2927 times (1510 times in the plus strand and 1417 times in the minus strand) in the *P. palmivora* genome. *In silico* PCR revealed 1672 fragments whose sizes varied from 175 bp to 3952 bp. The primer BLAST results in only a maximum of 1000 fragments for any set of primers.

### PCR assay for Mycobacterium tuberculosis

The IRS-based primer pair will amplify genomic regions randomly depending on positions that are complementary to the primer sequence in three possible priming combinations. Unlike conventional PCR, a pair of primers is not required in the IRS-based DNA amplification process. An IRS primer pair can amplify DNA via three different priming methods: i) forward primer alone, ii) reverse primer alone, and iii) forward-reverse combination. The genome mining algorithm identified identical repeat sequences from the *M. tuberculosis* genome amplified via the conventional PCR assay. Figure 2 shows that the *M. tuberculosis-*specific MtF1 (5′-CGC CGT TGC CGC CGT TGC-3′) and MtR1 (5′-GCC GCT CCT CCT CAT CGC-3′) primers alone amplified the DNA segments. On the basis of the gradient PCR results, we ran a PCR with a 58°C annealing temperature. These results were consistent with our expectation that the multiple primer binding locations of *M. tuberculosis* with short identical repeats would result in several amplicons. The gradient PCR of MtF1-MtR1 in combination resulted in multiple nonhomologous amplicons of different sizes (Figure 3). The targeting of multiple loci in a genome and the amplification of multiple copies improve the sensitivity and specificity of PCR-based diagnostic assays.

**Figure 2.**
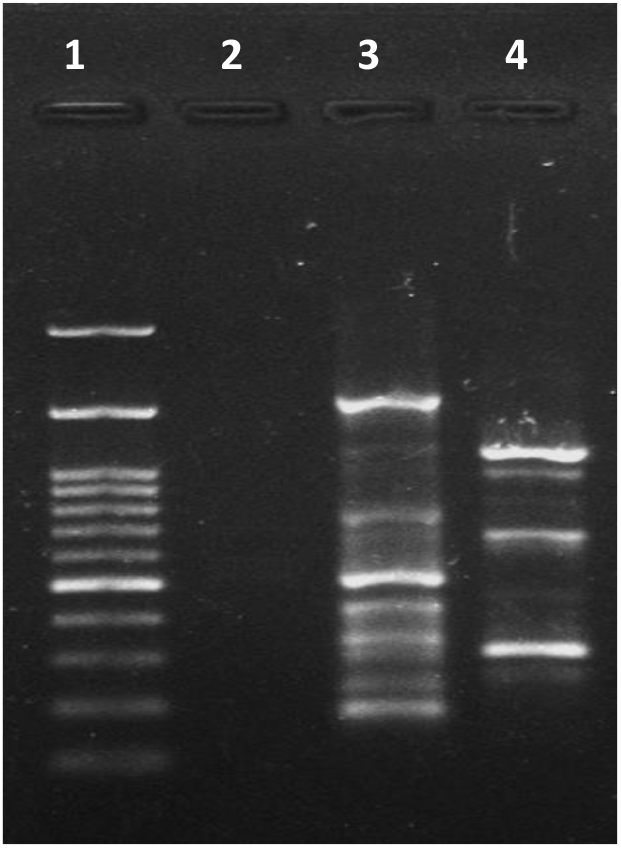
*M. tuberculosis* DNA fragments amplified via PCR with the individual primers MtF1 and MtR1 were run on a 1% agarose gel. Lane 1: 100 bp DNA ladder; lane 2: negative control; lane 3: PCR products of MtF1; lane 4: PCR products of MtR1.

**Figure 3.**
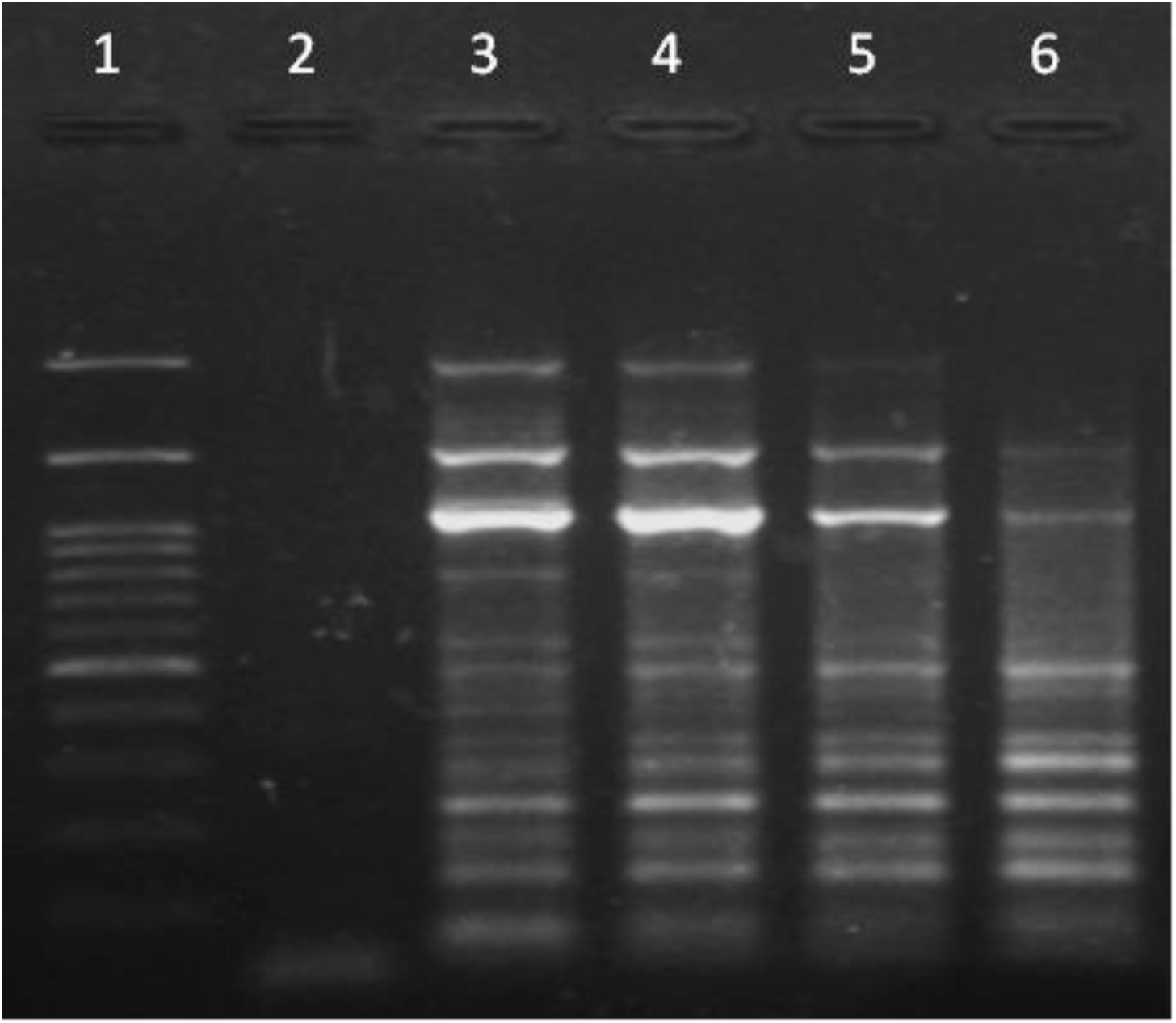
*M. tuberculosis* DNA fragments amplified via PCR with the pair of primers MtF1 and MtR1 were run on a 1% agarose gel. Lane 1: 100 bp DNA ladder; lane 2: negative control; lane 3: PCR products of MtF1-MtR1 annealed at 58°C; lane 4: PCR products of MtF1-MtR1 annealed at 60°C; lane 5: PCR products of MtF1-MtR1 annealed at 62°C; lane 6: PCR products of MtF1-MtR1 annealed at 64°C.

## Discussion

Currently, PCR-based approaches are extensively used to diagnose infectious diseases (Yang and Rothman, 2004). PCR-based tests are often highly sensitive and specific, and primers can be designed to detect a specific family, genus, or species without detecting closely related pathogens (van Weezep et al., 2019). Nucleic acid amplification or hybridization techniques offer direct detection of viral and bacterial pathogens from clinical samples (Johny et al., 2022). However, the success rates of DNA amplification or hybridization techniques are highly dependent on short primer or probe sequences. The use of primers or probes to initiate amplification or hybridization at multiple loci across the genome increases the analytical sensitivity of the currently used diagnostic assays. On the basis of primer targeting, DNA amplification mainly categorized into three main groups (Kalle et al., 2014). The first group of primers targets and amplifies a single type of DNA molecule via a primer pair. The second group contains a mixture of primers that target several nonhomologous regions and amplify different sequences simultaneously. The third group contains a single set of primers that target similar genomic regions and amplify a mixture of homologous DNA sequences. Our *in silico* and *in vitro* findings show that random amplification by a pair of IRSs in three priming combinations yields multiple amplicons in the PCR assay. Hence, the present study proposes a fourth group of primers, which can target short identical regions dispersed across the genome and amplify several nonhomologous regions with different fragment sizes in PCR assays.

### Algorithm for IMR identification

Over the years, several automated algorithms have been reported for identifying repeat sequences from genome sequences (Benson, 1999; Shi and Liang, 2019). A majority of these algorithms are efficient at identifying short tandem repeats but not for identifying repeats that can be used as primers or probes. One study developed an algorithm to identify the distribution of multiple identical repeat sequences as primers in the *Plasmodium falciparum* 3D7 genome and identified forward primers that were repeated 170 times and reverse primers that were repeated 123 times (Raju et al., 2019). We analysed the *P. falciparum* 3D7 genome with our algorithm to identify identical repeats. Interestingly, the algorithm identified forward IRS repeated 329 times and reverse IRS repeated 309 times. The repeat number is the total number of IRSs found on the plus and minus strands. *In silico* PCR of the IRS pair with the *P. falciparum* genome revealed 878 amplicons, indicating that primers derived from IRS are much more efficient at amplifying multiple copies of nonhomologous genomic regions. Thus, diagnostic assays with IRS can improve the overall sensitivity even with a low quantity of template DNA or pathogen in the incubation period.

A recent study developed degenerate primers that are capable of amplifying the conserved genes DPOL and gB at the lowest DNA dilution level for alpha, beta, and gamma herpesviruses that infect both humans and animals (Okoh et al., 2023). The PCR assay produces a bright single band that can be used for further sequencing or the identification of genotypes or other known herpesviruses. We identified the IRS from human gammaherpesvirus (Acc. No. LR994540.1), and *in-silico* PCR yielded 26 fragments without mismatches in the primer-binding region, varying in size from 36 to 1665 bp. *In-silico* PCR with default parameters revealed an additional 53 fragments with mismatches in the primer binding site on the template. These results suggest that the conversion of variable sites with appropriate degeneracy can yield more fragments and improve the overall assay sensitivity.

Short identical repetitive sequences have several advantages over gene- or multigene-specific primers or probes in diagnostic assays. Few studies have reported genome mining algorithms to improve the sensitivity of diagnostic assays. Most of these algorithms identify identical homologous regions dispersed throughout the genome, which amplify multiple genomic regions and thus improve sensitivity (Hofmann et al., 2015; Nolasco et al., 2021). Another set of algorithms involves identifying the conserved region present in the multigene converted as a primer and thus amplifying multiple genes in a single PCR assay (Schenk et al., 2017). The algorithm presented here will identify short IRS regions with primer or probe potential. On the basis of diagnostic interest, the algorithm is flexible for designing primers or probes for assays such as polymerase chain reactions, microarrays, loop-mediated isothermal amplification, and fluorescence *in situ* hybridization.

### Primer selection criterion

The genome mining algorithm identifies IRSs with user-defined lengths (k-mers) from any genome of interest. The algorithm provides three parameters for user inputs before executing the algorithm: i) repeat length, ii) minimum repeat count, and iii) minimum GC content. These parameters can be effectively used to obtain the IRS with primer or probe potential. The algorithm searches only for the IRS with the exact repeat length provided by the user. However, the algorithm extends the search for the parameters repeat count and GC content equal to or above the numerical value provided by the user. Thus, the algorithm identifies only the user-defined length identical sequences with the occurrence is equal to or greater than the user-defined repeat count and GC content. These parameters are crucial for researchers to identify identical repeats that are optimal for their diagnostic assay. A lower number (one or two digits) suggested that the genomes are small in size (<=1 Mb). An ideal primer contains 40–60% GC content and is suggested for the 3′ end with either G or C (Chuang et al., 2013). Some genomes are rich with AT/GC content, which reflects the short identical repeat sequence, and it may be difficult to identify the IRS with good primer characteristics. Suggested for a pair of IRSs (one each from the plus and minus strands) for PCR-based assays. Users must select the potential identical sequence with the optimum GC content for primer synthesis. Furthermore, the primer pair must have more or less the same melting temperature and should not have secondary structures. Available sophisticated computational tools can be utilized to assess the quality of the selected primers. Genomes may contain nested overlaps of short identical repeats or multiple homologous regions with single nucleotide polymorphisms. The algorithm identifies these genetic features uniquely and thus results in multiple repeats with one or two base differences between them. We provided a shell script to identify the exact number of identical sequences repeated without nested overlap present in a genome. A phylogenetic tree can be constructed to identify similar sequence clusters and develop degenerate primers.

No fragment will be amplified if the IRS acts as a forward or reverse primer or if the 3’ ends of the primer do not face each other. As each strain is unique, there will not be any amplification if the primer binding regions are not present on the template. Furthermore, mutation is quite common in microbial populations. Therefore, if there are mutations in the primer binding region on the template DNA, a PCR product will not be produced. Significant differences in the melting temperature of a primer pair may make it difficult to set an appropriate annealing temperature and may require gradient PCR. Furthermore, the IRS of mononucleotide repeats can be avoided. An identical sequence repeated at a maximum in the genome may not amplify DNA fragments. The IRS 3’ ends facing each other with maximum loci less than 3 kb in distance yield more amplicons via conventional PCR. Primer-BLAST can be effectively utilized to prioritize the best IRS, yield a maximum number of fragments and identify species-specific IRSs. Differences in genome sequencing technologies and genome assembly algorithms may result in different banding patterns of amplified DNA fragments on the gel compared with *in-silico* PCR results.

### Comparison of IRSs with multicopy genes and RAPD markers

The 16S or 18S rRNA genes are the most commonly used amplification targets of pathogens in clinical diagnosis, as they have highly conserved genera and species-specific regions and present five to eight copies per genome (Wang et al., 2014; Hofmann et al., 2015; Bartoš et al., 2024). However, the primers derived from IRS can amplify multiple fragments, as they can amplify randomly. Thus, the IRS can provide better sensitivity than multigene amplification assays. Furthermore, several reports on the high interspecies identity of 16S rRNA have been published (Beye et al., 2017), and 18S rRNA rarely resolves fungal taxa at the species or genus level (Yarza et al., 2017). Genus- or species-specific primers or probes can be developed from the IRS, as the algorithm delivers several short patterns. Random amplified polymorphic DNA (RAPD) markers are simple and cost-effective, and no prior knowledge of the genome under study is necessary. It can also be performed in small laboratories for most applications (Babu et al., 2021). The algorithm we developed requires a whole-genome sequence to identify the IRS. However, the individual IRS can amplify multiple copies and even more copies in pairs. Overall, the IRS primer can yield better amplification than other currently available approaches.

## Limitations

Several studies have reported that misassembly can even persist in both draft and finished genomes and that these assemblies may mislead further genome-based studies. Forward (plus) and reverse (minus) stranded genomic regions can be found on a single strand of an assembled genome (Phillippy et al., 2008; Greenberg and Shomorony, 2022). Hence, the genome assembly used for the discovery of IRSs with forward- and reverse-strand breakpoints on the same strand may not be amplified in PCR assays. The strand orientation may differ in the template DNA used for PCR amplification. The reverse complement of the IRS pair may be amplified in such cases. Furthermore, *in silico* PCR amplifies genomic regions in all possible priming combinations, including the IRS, in a nested repeat format. However, all the nested repeats may not be amplified in conventional PCR assays.

## Conclusion

In summary, the algorithm described herein can efficiently identify short identical repeat sequences with a user-defined k-mer length. This algorithm can be used as a platform technology for developing genome-based approaches for various diagnostic assays. Using this algorithm, we identified identical repeat sequences from viruses, bacteria, parasites, and oomycetes. Furthermore, the identical repeat sequences identified from *Mycobacterium tuberculosis* revealed several amplicons via conventional polymerase chain reaction. Moreover, the algorithm is flexible enough to identify potential primers and probes for DNA amplification or hybridization assays. The identical repeat sequences that we identified can be utilized for the development of diagnostic assays for various diseases and genome-based approaches for species identification. We believe that molecular diagnostic assays incorporating identical repeat sequences will provide high sensitivity, as they can yield multiple nonhomologous amplicons of variable sizes via three possible primes. Therefore, the genome mining algorithm for identical repeat-based biomarker discovery translates infectious disease diagnostic approaches in a better manner.

## Acknowledgements

BP would like to thank Manipal Academy of Higher Education (MAHE), and the Department of Biotechnology (DBT-BUILDER scheme), Government of India for the infrastructure and support.

## Notes

### Competing Interest Statement

The authors have declared no competing interest.

